# Effects of prey abundance and foliage structure in plant selection by insectivorous birds in the central Monte desert

**DOI:** 10.1101/2024.12.11.627985

**Authors:** Carolina Guerra-Navarro, Victor R. Cueto

## Abstract

Foraging behavior allows a direct examination of how birds use their habitats and provides the possibility to understand what environment factors can determine those patterns. We analyzed the woody plant selection, height and substrate use, and attack maneuver by Greater Wagtail-tyrant (*Stigmatura budytoides*), Grey-crowned Tyrannulet (*Serpophaga griseicapilla*) and Ringed Warbling-Finch (*Microspingus torquata*) in the main habitats of central Monte desert. The three bird species selected *Prosopis flexuosa* trees and avoided the most abundant shrub species: *Larrea divaricata* and *Larrea cuneifolia* in mesquite woodland and creosotebush habitats. The importance of *P. flexuosa* was due because this plant constitute an abundant food patch of great diversity of arthropods. Our results about how bird species use their habitat and which factors were influencing these selection pattern are important to management and conservation of insectivorous birds in the central Monte desert.

## Introduction

Studies on foraging behavior allows a direct examination of how birds use their habitats and provides the possibility to understand what environment factors can determine those patterns (Holmes et al. 1979). The woody plant species selection as foraging sites is a well-documented phenomenon with regard to bird foraging behavior (e.g., Holmes and Robinson 1981, Greenberg and Bichier 2005, Cueto and Lopez de Casenave 2002, Gabbe et al. 2002, Oyugi et al. 2012). Several studies have found that the type and abundance of preys could determine the selection of a plant species by birds (Holmes and Robinson 1981, Holmes and Schultz 1988, Whelan 1989, Robinson and Holmes 1984, Hino et al. 2002). However others factors as the foliage structure could besides determine the bird plant selection (Holmes and Robinson 1981). Several studies have found that foliage structure of different plant species provide different foraging opportunities and constraints that determine how and where birds forage (Robinson and Holmes 1982, Holmes 1990). Both field (Holmes and Robinson 1981, Unno 2002, Park 2005, Park et al. 2008) and experimental studies (Parrish 1995, Whelan 1989, 2001), suggest that birds select plant species with foliage structure that facilitates prey encounter and capture. For example, gleaning birds usually select plant species with dense foliage, small leaves and short petioles (Holmes and Robinson 1981, Robinson and Holmes 1982, 1984, Cueto and Lopez de Casenave 2002), while hovering birds are less affected by differences in foliage structure and do not show strong selection for particular tree species (Holmes and Robinson 1981). Also, bird species can modify their foraging behavior between habitats with different structural complexity of vegetation (Maurer and Whitmore 1981). Several studies have found that birds changed tree species use (Maurer and Whitmore 1981, Block 1990), foraging heights (Maurer and Whitmore 1981), maneuvers use (Lyons 2005) and foraging substrate use (Tebbich et al. 2004) when these variables are analyzed in different habitat types.

The bird community of the central Monte desert is structured in several guilds, being the foliage-foraging insectivorous one of the most important (Lopez de Casenave et al. 2008). We examined the foraging behavior and microhabitat use of three insectivorous bird species from that guild during the breeding season: Greater Wagtail-tyrant (*Stigmatura budytoides*), Grey-crowned Tyrannulet (*Serpophaga griseicapilla*) and Ringed Warbling-Finch (*Microspingus torquata*). We analyzed the woody plant selection, height and substrate use, and attack maneuver by birds in the woody plant species of in the main habitats of central Monte desert: mesquite open woodland and creosotebush. We examined the following factors in the woody plant species: (1) abundance and biomass of arthropods in the foliage and (2) fine-scale foliage structures. We expect that if plant selection depends on the abundance of prey, the birds will select the woody species with the greater abundance or biomass of arthropods. Otherwise, if the plant selection depends on the foliage structure, the birds will be select the plants that have foliage that facilitates prey encounter and capture according their capture maneuvers.

## Methods

### Study area

The study was carried out in the Biosphere Reserve of Ñacuñán (34° 03’ S, 67° 54’ 30’’ W), located in the central Monte desert, Mendoza, Argentina. The landscape of the reserve is mainly a mesquite open woodland intersected by variably sized tracts of creosotebush shrubland. The mesquite woodland is a matrix of non-thorny tall shrubs (mainly *Larrea divaricata*), with thorny trees (*Prosopis flexuosa* and *Geoffroea decorticans*) and tall shrubs (*Capparis atamisquea* and *Condalia microphylla*). Nonthorny tall shrubs (*Larrea cuneifolia*) dominate the creosotebush shrubland, with scarce cover of thorny trees and tall shrubs (Marone 1991). The open woodland has higher vertical complexity (three vegetation strata) than the shrubland (two strata) (Marone 1991). The climate is dry and highly seasonal. Mean annual rainfall is 342 mm, and more than 75% of the annual rainfall occurs in the warmer months (October–March). Therefore, the summer and spring are hot and rainy and the winter and autumn are cold and dry.

Foraging behavior was studied on three plots of 10-ha (200 x 500 m) each one: two of them placed within mesquite woodlands and the third in a creosotebush shrubland.

### Vegetation sampling

We quantified the horizontal and vertical vegetation cover in the three plots. Each plot was divided into squares of 25 m x 25 m and the corners were marked with color flagging. We used a thin aluminum pole erected (graduated at 25 cm intervals) at 189 random points within each plot. Each point was determined with a random orientation and distance (between 0 and 12 m) from each of the square’s corners. At each point we recorded the plant species that touched the pole at each 25 cm interval up to 6,5 m. Horizontal cover for every species was estimated as the percentage of the total sampled points where each species was recorded. We also made a height profile of foliage cover as the percentage of points with contacts at 1- m interval.

### Bird foraging behavior

We recorded the foraging behavior of Greater Wagtail-tyrant (*Stigmatura budytoides;* 11 g) and Grey-crowned Tyrannulet (*Serpophaga griseicapilla;* 5.5 g) belonging to the family Tyrannidae, and Ringed Warbling-Finch (*Microspingus torquata;* 10 g) belonging to the family Thraupidae (this latter species shifts from the foliage-foraging guild to the graminivore guild during the non-breeding season, Lopez de Casenave et al. 2008). Greater Wagtail-tyrant and Ringed Warbling-Finch are resident birds in the study area (Marone 1992); Grey-crowned Tyrannulet is a short migrant bird, present at the study area during spring and summer (Cueto et. al 2008).

We gathered foraging data during three breeding-seasons: 2006-2007, 2007-2008 and 2008-2009. During each sampling period, we systematically walked through the three plots from sunrise to midday and from the afternoon until sunset, except on rainy or too windy days. We avoid walked through the same area twice on the same day to avoid repeating the observations of the same individual birds. Each time we observed a bird searching for prey, we followed it as long as we could keep it in sight, and we registered the following variables in a portable digital voice recorder: bird species, woody plant species, substrate from which food was obtained, prey-attacking maneuvers and height of attack estimated to the nearest 1 m. Substrates considered were: branches, twigs, foliage (including leaves and petiole), gall, flowers and air.

Prey-attacking maneuvers were sorted into three categories (based on Remsen and Robinson, 1990): (1) glean: when a perched bird captures food from the surface of a nearby substrate; (2) sally-hover: when a bird captures food from the surface of a substrate while in flight; (3) sally-strike: when a bird flies from a substrate to capture a prey in the air.

Data from the three breeding seasons were pooled for each bird species to analyze the foraging behavior. We used the sequential observations of each individual bird to evaluate the foraging behavior because the initial observations can be subjected to a conspicuousness bias (since the birds can be more visible at some parts of their habitat, Morse 1990). Moreover, the sequential observations might reveal more about the foraging behavior of these species than the initial observation (Morse 1990). However the sequential observations are not independent of one another (Hejl et al. 1990, Morse 1990). In order to avoid this problem we used each sequence of observations from an individual as a single observation when we determined frequencies and sample sizes for statistical tests (Airola and Barret 1985). Thus, when *n* consecutive observations of an individual were recorded, each observation contributed to the species’ total frequency by a value of 1/*n,* and all the observations from the same individual contributed *Σ1 /n = 1* to the species’ frequencies (Airola and Barrett 1985).

We evaluated woody plant species selection with the Chi-square goodness-of-fit test for each plot study. We tested the hypothesis that observed distribution of attacks among woody plant species for each bird species was the same as the expected distribution obtained from the relative cover of woody plant species. We also evaluated foraging height use with the Chi-square goodness-of-fit test. We separated foraging height data into 1 m height categories. To the woody plants selection and foraging height we only considered glean and sally-hover attacks because these maneuvers are carried out by the birds directly to the woody species, unlike the sally-strike that are not foliage-related.

To achieve adequate expected cell values for the chi-square tests, we combined adjacent categories in some cases (of woody plant species or height depending on the analysis) to raise the expected values following the less restrictive guidelines for chi-square goodness of fit testing of Roscoe and Byars (1971), carrying the average expected frequency for all cells be greater than 4 and no cell should have an expected frequency lower than 1. A consequence of that procedure was the reduction of degrees of freedom in some cases.

We compared the foraging maneuvers and substrates of each bird species in the three plots. We tested also the independence between the plant species and the foraging maneuvers realized by the birds with Chi-square test (sally was not included in this analysis). For this analysis we pooled the data of the three plots for each bird species to get a great number of attack observations for each plant species since there were no differences in the maneuvers pattern between the two habitat types.

### Arthropods sampling

We used the “branch clipping” method (Cooper and Whitmore 1990, Johnson 2000) to evaluate the arthropod abundance and biomass in the plants. The arthropods sampling was carried out in the same period of the observations of birds foraging behavior. We sampled arthropods on six woody species (*Prosopis flexuosa, Geoffroea decorticans, Capparis atamisquea, Condalia microphylla, Larrea divaricata* and *L. cuneifolia*) in each sampling period (spring and summer of each three breeding seasons). The sampling was carried out between 10:30 am to 1:30 pm. We established ten sampling stations at one mesquite woodland plot and ten at the creosotebush shrubland plot in which we randomly selected one individual of each plant species and measured: total height, height from the ground to the first branch with foliage, longest canopy diameter and perpendicular diameter to the longest. We collected an arthropods sample enclosing one branch (approximately 50 cm) into a plastic bag as quickly as possible. Then we clipped the branch with shears and fumigated the sample with insecticide. For each sampled branch we recorded its length, the longest branch diameter and the perpendicular diameter to the longest. We quantified the number of arthropods recovered from each sampled branch up, these arthropods were identified to Order or Family level and their body length was measured with a micrometer under a binocular microscope; we only considered the arthropods larger than 1 mm following others studies (Wiens et al. 1991, Diaz et al. 1998). We calculated the diversity of arthropods sorted by taxa (orders and families) with the Shannon’s diversity index: H= - Σ(p_i_) (ln p_i_), where p_i_ is the proportion of arthropods belonging to category i (order/family).

The dry weight (W, mg: 60°C, 48 h) of arthropod was estimated from the body length (L, mm) with equations derived from weights and lengths measured on a subset of the samples collected at the study site (following Hódar 1996). We used different equations based in the order/family level and we taked into account morphological characteristics as shape and body toughness (Hódar 1996). We carried out in total six equations (see Apendix 1).

For each sample we estimated the volume for the clipped branch and the whole plant, assuming shapes of different geometrical figures for each one. We considered all branches as cylinder-shaped, and for the whole plants we considered the canopy of *P. flexuosa*, *C. microphylla* and *C. atamisquea* as a hemisphere; *L. cuneifolia* and *L. divaricata* as a cone; and *G. decorticans* as a cylinder. We then estimated the abundance and biomass of arthropods per m^3^ and per volume of the individual plant. We pooled the arthropod data of the three breeding seasons for all the analyses. With the data from 5 variables obtained in each arthropod sampled (Abundance/m^3^, Abundance per individual plant, Biomass/m^3^, Biomass per individual plant, Diversity index) from the six plant species at the two habitats types we carried out a Principal Component Analysis, through which reduced the number of variables in a few, clearly identifiable, principal components (axes) without lost of information.

### Foliage structure

To quantify fine-scale foliage structure we used a modification of the methods used by Whelan (2001) and Park et al. (2008). The measurements recorded in the foliage plants help us to classify foliage structure according to the degree of dispersion and density of branches and leaves; these characteristics (e.g. distance to nearest branch, petiole length, leaf area and leaf dispersion around supporting branches) can affect bird foraging behavior (Holmes and Robinson 1981, Whelan 2001). We measured the foliage variables in 10 random individuals of each of the following plant species: *P. flexuosa, G. decorticans, L. divaricata, L. cuneifolia, C. atamisquea* and *C. microphylla*. To evaluate the branch dispersion we measured the average distance to the nearest four branches in two branches of each individual (in the middle and outside region of the plant individual). To evaluate the leaves dispersion we cut off a 50 cm branch from each plant individual; we quantified the number of leaves on three 10 cm sections located at the inner, middle and distal portion of the collected branch (including leaves attached to the target branch as well as leaves attached to other branches within a radius of 5 cm of the target branch). We also took digital pictures from 10 leaves of each individual plant and measured leaf area using Scion Image program. We also classified 30 random leaves of each plant species according to its position respected to the plane horizontal to the twig: above, even with, or below. We measured the petiole-length of 30 random leaves from the branch collected of each individual. With the data from the 9 variables measured in each of the ten individuals from the six plant species we set a 60 x 9 matrix and subjected it to a Principal Component Analysis.

## Results

### Plant species selection

The three plots had different percentage cover of woody plant species, especially *P. flexuosa* that had more than double of cover in the mesquite woodland than creosotebush shrubland (Table 1). The three bird species used *P. flexuosa* as feeding patch more frequently than expected from its relative abundance in the three plots (Table 1). The most abundant woody plant species in each of the two habitat types (*L. divaricata* in mesquite woodlands and *L. cuneifolia* in creosotebush shrubland) was avoided by birds (Table 1). All others woody plant species were used according to their availability (Table 1).

**Table 1.** Cover percentage of woody plant species en the three studied plots (two mesquite woodlands and one creosotebush shrubland at Ñacuñán Reserve, Mendoza) and percentages of use by birds. Chi-square goodness of fit comparison between plant availability and use are shown for each bird species and each study plot. Code of woody species: Pf: *Prosopis flexuosa*; Ld: *Larrea divaricata*, Gd: *Geoffroea decorticans*, Ca: *Capparis atamisquea*, Cm: *Condalia microphylla* and Lc: *Larrea cuneifolia*.

|  | Woody plant species |  |  |  |  |  |  |  |  |
| --- | --- | --- | --- | --- | --- | --- | --- | --- | --- |
| | Pf | Ld | Gd | Ca | Cm | Lc | N | $X^2_{df}$ | P |
| **<u>Mesquite woodland 1</u>** |  |  |  |  |  |  |  |  |  |
| **Cover (%)** | 12.2 | 43.9 | 3.7 | 1.6 | 2.1 | 4.2 |  |  |  |
| **Plants use by birds (%)** |  |  |  |  |  |  |  |  |  |
| Greater Wagtail-tyrant | 59.8 | 14.3 | 8.6 | 8.0 | 4.7 | 4.7 | 67 | $X^2_5= 101.9$ | <0.001 |
| Grey-crowned Tyrannulet | 84.3 | 10.9 | 3.9 | 0.4 | 0.6 | - | 90 | $X^2_5= 269.4$ | <0.001 |
| Ringed Warbling-Finch | 83.4 | 4.8 | 5.1 | 4.1 | 1.7 | 0.8 | 118 | $X^2_5= 353.4$ | <0.001 |
| **<u>Mesquite woodland 2</u>** |  |  |  |  |  |  |  |  |  |
| **Cover (%)** | 23.8 | 37 | 7.9 | 3.7 | 2.1 | 0.5 |  |  |  |
| **Plants use by birds (%)** |  |  |  |  |  |  |  |  |  |
| Greater Wagtail-tyrant | 68.1 | 14.4 | 11.6 | 5.8 | - | - | 60 | $X^2_4= 42.0$ | <0.001 |
| Grey-crowned Tyrannulet | 78.5 | 9.6 | 10.2 | - | 1.7 | - | 59 | $X^2_4= 63.1$ | <0.001 |
| Ringed Warbling-Finch | 80.3 | 2.1 | 12.8 | 4.8 | - | - | 47 | $X^2_4= 58.0$ | <0.001 |
| **<u>Creosotebush shrubland</u>** |  |  |  |  |  |  |  |  |  |
| **Cover (%)** | 5.8 | 14.3 | 2.6 | 1.1 | 3.2 | 45.5 |  |  |  |
| **Plants use by birds (%)** |  |  |  |  |  |  |  |  |  |
| Greater Wagtail-tyrant | 77.7 | - | 2.3 | 10.7 | 2.3 | 7.0 | 43 | $X^2_4= 294.6$ | <0.001 |
| Grey-crowned Tyrannulet | 50.7 | 10.2 | 19.4 | - | - | 19.6 | 18 | $X^2_3= 49.1$ | <0.001 |
| Ringed Warbling-Finch | 74.7 | - | 5.2 | 12.1 | 3.4 | 4.7 | 58 | $X^2_4= 374.1$ | <0.001 |

### Foraging maneuvers

Each bird species showed a different pattern of foraging maneuvers, but they did not changes their maneuvers attacks at the three different plots (Fig. 1). Greater Wagtail-tyrant and Ringed Warbling-Finch captured their prey mainly by glean, while Grey-crowned Tyrannulent most frequently use sally-hovered (Fig. 1). The frequency of gleaning and sally-hovering was independent to the woody plants for the three bird species (Table 2).

**Fig. 1.**
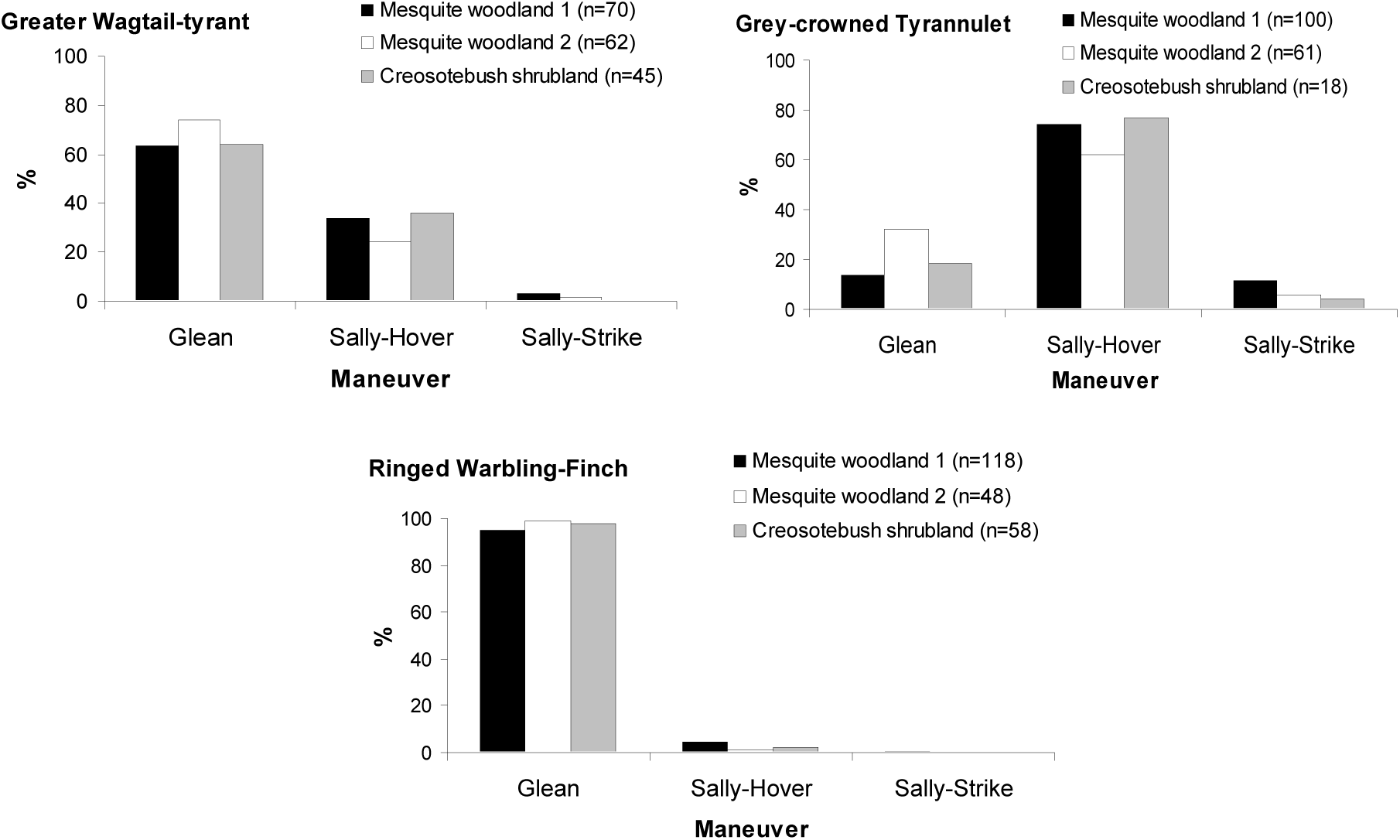
Percentage of foraging maneuvers used by the three bird species in two mesquite woodlands and one creosotebush shrubland of Ñacuñán Reserve during three breeding seasons.

**Table 2.** Percentage of foraging maneuvers used by three bird species on each woody plant species. Data was pooled from observations at the mesquite woodland and the creosotebush shrubland of Ñacuñán Reserve during three breeding seasons. Independence between the foraging maneuver and woody plant species was evaluated by Chi-square testing: sample size (N), Chi-square value (X^2^), and *P*-value (*P*) are shown for every test. See Table 1 for plant species code.

| Use of maneuver by birds (%) | Woody plant species | | | | | | N | $X^2_{df}$ | P |
| --- | --- | --- | --- | --- | --- | --- | --- | --- | --- |
|  | Pf | Ld | Gd | Ca | Cm | Lc |  |  |  |
| **Greater Wagtail-tyrant** | | | | | | | 170 | $X^2_5 = 2.47$ | 0.78 |
| Glean | 66.3 | 66.8 | 83.0 | 78.9 | 75.8 | 63.4 |  |  |  |
| Hover | 33.7 | 33.2 | 17.0 | 21.1 | 24.2 | 36.6 |  |  |  |
| **Grey-crowned Tyrannulet <sup>1</sup>** | | | | | | | 167 | $X^2_2 = 5.36$ | 0.07 |
| Glean | 19.4 | 31.8 | 43.6 | 100.0 | 33.3 | 28.3 |  |  |  |
| Hover | 80.6 | 68.2 | 56.4 | 0.0 | 66.7 | 71.7 |  |  |  |
| **Ringed Warbling-Finch <sup>2</sup>** | | | | | | | 222 | $X^2_1 = 0.25$ | 0.62 |
| Glean | 96.1 | 97.5 | 100.0 | 98.6 | 100.0 | 92.6 |  |  |  |
| Hover | 3.9 | 2.5 | 0.0 | 1.4 | 0.0 | 7.4 |  |  |  |

<sup>1-2</sup> Woody plants categories were combined to make expected frequencies large enough for chi-square test.
<sup>1</sup> Three woody plants species categories: Pf, Ld+Lc, Gd+Ca+Cm
<sup>2</sup> Two woody plants species categories: Pf, Ld+Ca+Cm+Gd+Lc

### Foraging substrates and height distribution

The three species concentrated their foraging mainly on foliage (Fig. 2). Twigs also were used, but in low frequencies. Ringed Warbling-Finch used flowers more than the other two species, and Grey-crowned Tyrannulet frequently captured prey in the air more than the other two species (Fig. 2).

**Figure 2.**
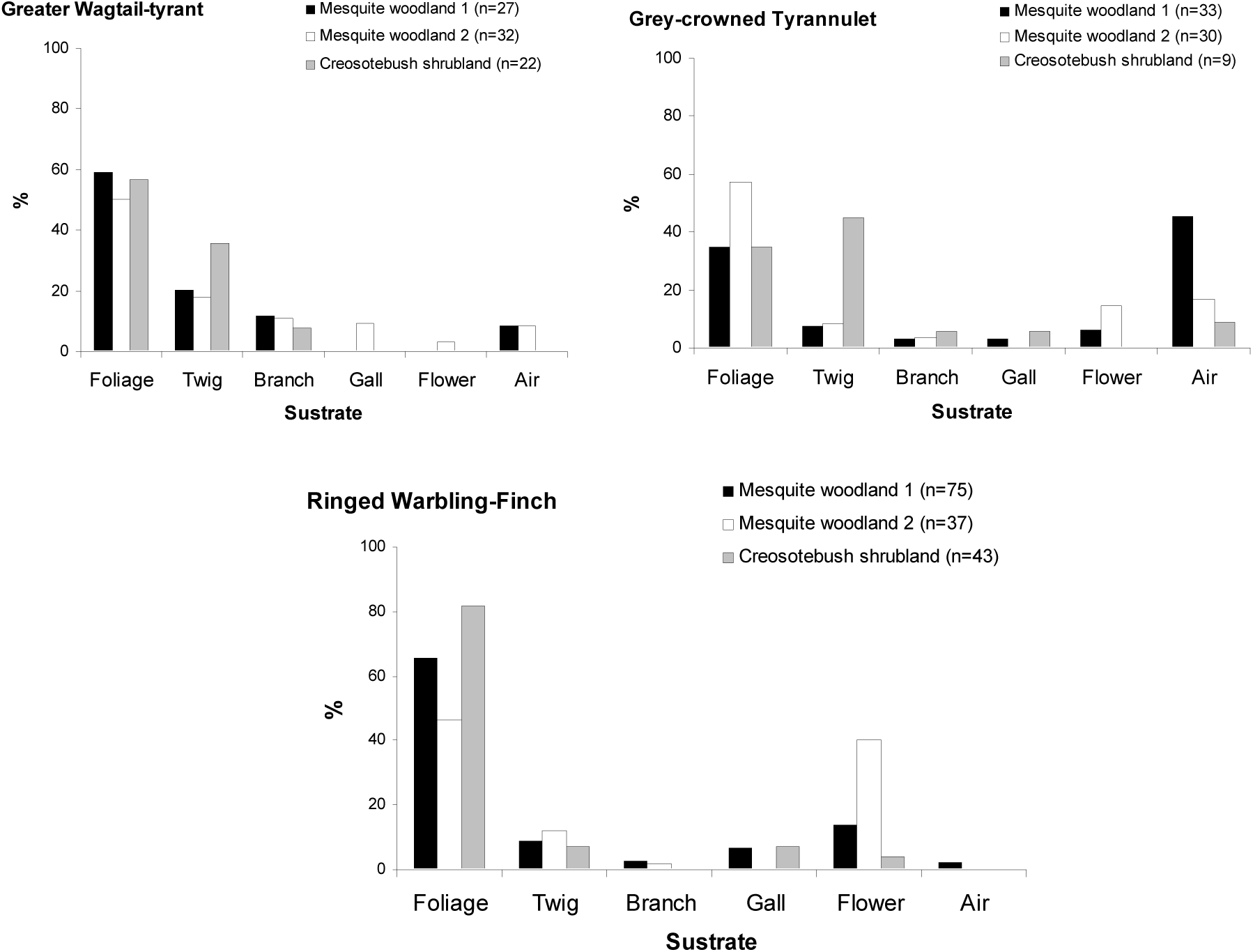
Percentage of substrate used to forage by the three bird species in two mesquite woodlands and one creosotebush shrubland of Ñacuñán Reserve, Mendoza, during three breeding seasons.

The height pattern of available foliage of the woody plants differed between plots, mesquite woodlands have more cover than creosotebush shrubland in almost all height (Fig 3a). The three bird species captured prey predominantly in the range of height from 2 to 4.5 m (Greater Wagtail-tyrant: X^2^ = 70.9, *P* < 0.001; X^2^ =34.0, *P* < 0.001 and X^2^ =62.9, *P* < 0.001, Fig 3b; Grey-crowned Tyrannulet: X^2^ = 230.4, *P* < 0.001; X^2^ =62.2, *P* < 0.001 and X^2^ =39.1, *P* < 0.001, Fig 3c and Ringed Warbling-Finch: X^2^ = 189.3, *P* < 0.001; X^2^ =72.3, *P* < 0.001 and X^2^ =65.2, *P* < 0.001, Fig 3c).

**Figure 3:**
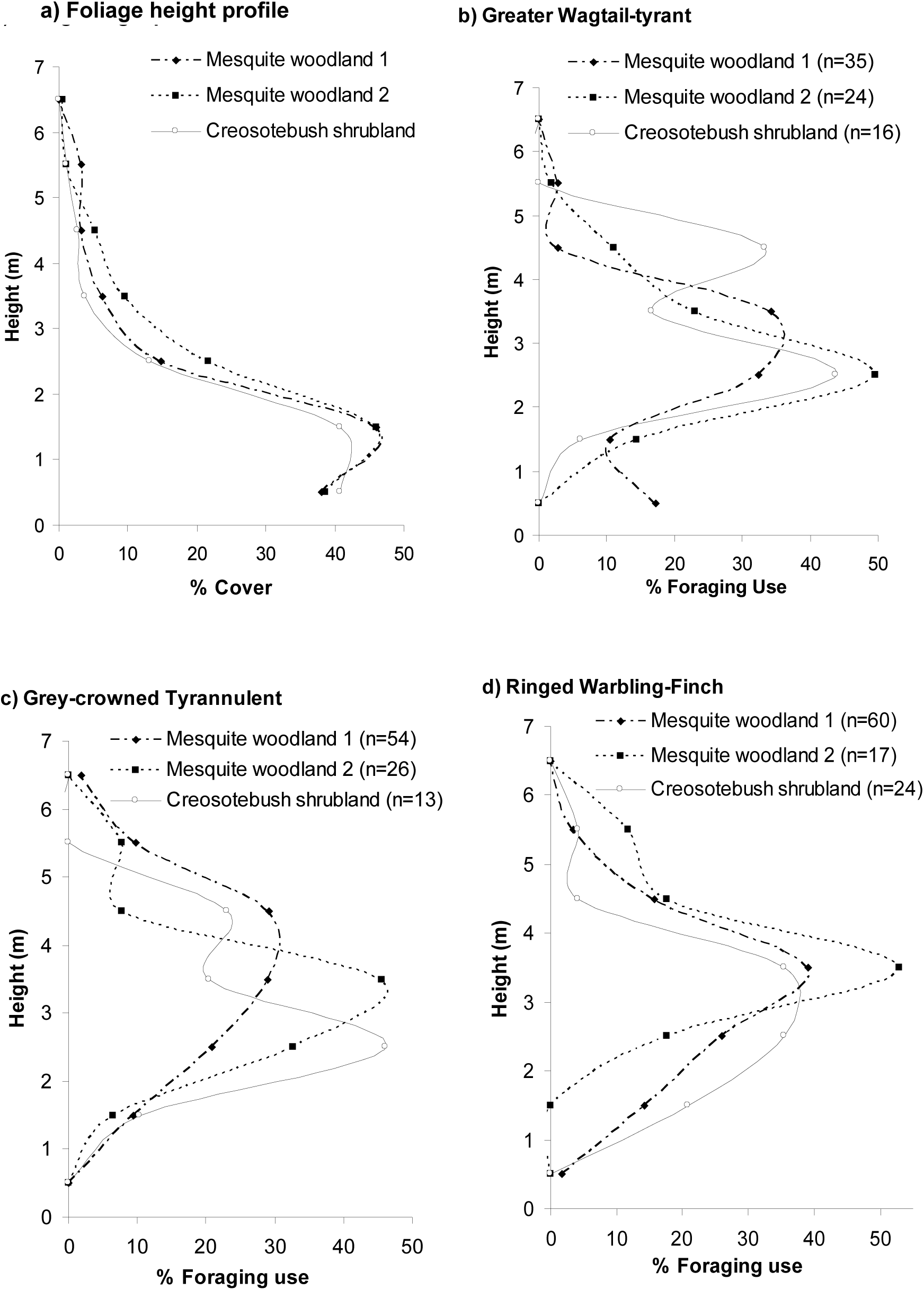
a) Foliage height profile and b-d) foraging height distribution of three bird species in three plots at Ñacuñán Reserve, Mendoza, during three breeding seasons.

### Arthropods

We collected 6321 arthropods in 600 branch samples. The Principal Components Analysis allowed us to discriminate the most important variables characterizing the arthropod samples of six plant species at the two habitats. The first three components accounted for 84.5 % of the total variance (Table 3). The first two components can be used to explore variations in the abundance of arthropods among the plant species. The first axis was positively associated with the abundance and biomass of arthropods in each plant individual (Table 3).This axis reflects a contrast between *P. flexuosa* with the other plant species; this species was located to the positive values of the first axis possessing the highest values of abundance and biomass per individual due to their higher canopy volume compared with the other plant species (Fig. 4, Table 4). The second component showed negative loadings for the diversity of orders and families of arthropods, with *P. flexuosa* with scores more negatives in this axis, and both *Larrea* spp. with positive values. So, the ordination separated to *P. flexuosa* of the rest of the plant species as the species with greater abundance and biomass of arthropods and with the greater Shannon’s diversity index of orders and families of arthropods (Fig. 4, Table 4).

**Table 3:** Principal Component Analysis based on arthropods variables of six woody plant species at two habitats from Ñacuñán Reserve, Mendoza. Loadings for the most heavily weighted variables are shown in bold.

| *Variables* | *Code* | *Components* |  |  |
| --- | --- | --- | --- | --- |
|  |  | CP 1 | CP 2 | CP 3 |
| Arthropods Abundance / m<sup>3</sup> | Abund m<sup>3</sup> | 0.57 | **0.69** | 0.03 |
| Arthropods Abundance / individual plant | Abund ind | **0.82** | -0.28 | 0.32 |
| Arthropods Biomass / m<sup>3</sup> | Biom m<sup>3</sup> | 0.54 | 0.44 | **-0.64** |
| Arthropods Biomass / individual plant | Biom ind | **0.79** | -0.37 | 0.19 |
| Arthropods Diversity | Divers | 0.15 | **-0.73** | -0.58 |
| Eigenvalue |  | 1.92 | 1.41 | 0.89 |
| % Variance |  | 38.4 | 28.2 | 17.9 |
| Cumulative |  | 38.4 | 66.6 | 84.5 |

**Table 4:** Prey abundance and biomass of arthropods per m^3^ and per individual plant ± SE of six woody plant species in two habitats in Ñacuñán Reserve during three breeding seasons. The number and biomass of arthropods per individual plant were estimated based in the different mean canopy volume of the different species (see Methods). See Table 1 for plant species codes. N= 60 for each plant species at mesquite woodland (“mesq”) and N= 40 at creosotebush shrubland (“creo”).

|  | N° Arthropods/<br>m<sup>3</sup> | Biomass (mg) / m<sup>3</sup> | Canopy<br>volume (m<sup>3</sup>) | N° Arthropods /<br>individual plant | Biomass (mg) /<br>individual plant | Diversity<br>index (H) |
| --- | --- | --- | --- | --- | --- | --- |
| Pf mesq | 502.9 ± 72.0 | 204.3 ± 34.7 | 203.6 ± 17.3 | 100912 ± 12855.9 | 53597.4 ± 10943 | 1.91 |
| Pf creo | 719.1 ± 153.6 | 156.8 ± 27 | 153.5 ± 14.3 | 108371.4 ± 18366.4 | 23475.1 ± 3018.9 | 1.82 |
| Ld mesq | 387.4 ± 56.9 | 138.9 ± 29.4 | 6.6 ± 0.5 | 2332.6 ± 232.2 | 940.6 ± 121.4 | 1.52 |
| Ld creo | 782 ± 171.2 | 129 ± 25.8 | 4.8 ± 0.5 | 3051.9 ± 502.7 | 541.5 ± 61.6 | 1.16 |
| Gd mesq | 229.3 ± 22.7 | 71.8 ± 9.6 | 7.6 ± 0.6 | 1814.2 ± 219.3 | 666.4 ± 87.6 | 1.62 |
| Gd creo | 203.8 ± 24.6 | 65.9 ± 13.7 | 13.9 ± 1.6 | 2898 ± 368.5 | 968.8 ± 152.8 | 1.54 |
| Ca mesq | 346.7 ± 35.5 | 200.5 ± 42.1 | 27.5 ± 2.7 | 10437.2 ± 1312.3 | 11793.7 ± 2374.8 | 1.7 |
| Ca creo | 391.7 ± 53.9 | 183.9 ± 36.0 | 35.7 ± 4.0 | 13459.5 ± 1704.1 | 5931.7 ± 657.7 | 1.72 |
| Cm mesq | 522.9 ± 68.1 | 173.6 ± 21.2 | 7 ± 0.7 | 3659.1 ± 525.0 | 1546.5 ± 260.6 | 1.85 |
| Cm creo | 798.9 ± 128.5 | 276.1 ± 46 | 7.9 ± 1.0 | 5989.7 ± 780.2 | 2590.4 ± 326.2 | 1.79 |
| Lc mesq | 486.7 ± 62.8 | 189.5 ± 49.1 | 7.1 ± 0.7 | 3564.1 ± 415.3 | 1858 ± 300.8 | 1.29 |
| Lc creo | 909.4 ± 114.3 | 230.2 ± 48.9 | 5.6 ± 0.5 | 4883.9 ± 600.7 | 1330 ± 147.5 | 0.88 |

**Fig. 4.**
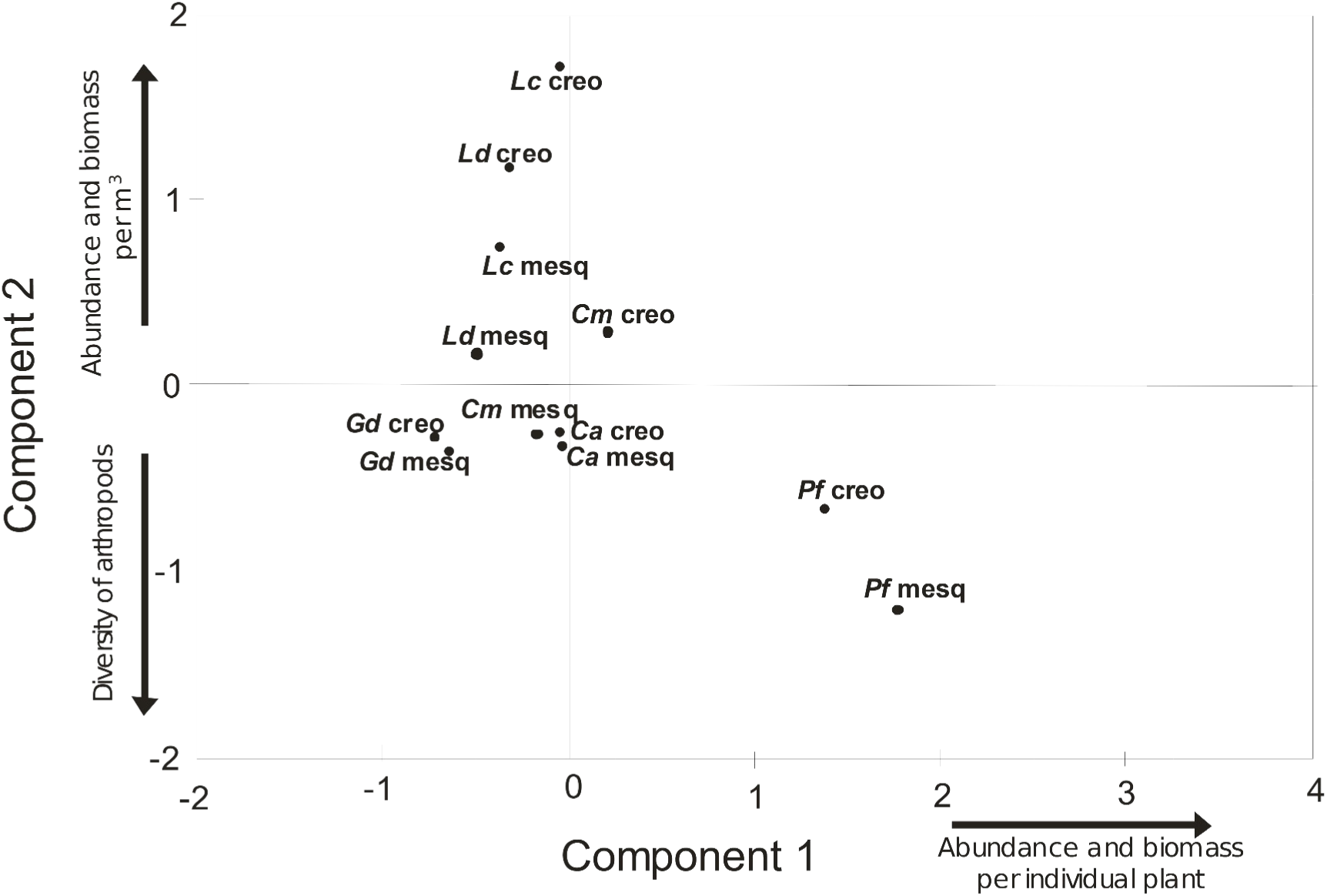
Position of woody plant species of two habitats from Reserve of Ñacuñán along the first two axes obtained from a Principal Component Analysis carried out with arthropods data. See Table 3 for loadings of variables for each component. See Table 1 for plant species codes. N= 60 for each plant species at mesquite woodland (“mesq”) and N= 40 at creosotebush shrubland (“creo”).

### Foliage structure

The Principal Components Analysis allowed us to discriminate the most important variables characterizing the foliage structure of the analyzed plants. The first three components accounted for 74.7 % of the total variance (Table 5). The first two components can be used to explore variations in the species’ foliage structure. The first axis was negatively associated with the number of leaves at the three position of the branch, and positively associated with petiole length, leaf area and the frequency of leaves below the plane of the branch (Table 5). This axis reflects a contrast between the plants with few and relatively large leaves per branch and large petioles (as *P. flexuosa* that are located to the positive values of the axis, Fig. 5), as opposed to plants with high number of little leaves with short petiole (as *C. microphylla* to the negative values of the axis) (Fig. 5). The second component showed negative loadings for number of leaves at the inner of the branch, with *C. microphylla* with scores more negatives in this axis, and both *Larrea* spp. with positive values. The ordination separated to *C. microphylla* as the species with foliage more clustered (high number of leaves, short petioles and a similar disposition of leaves along the branch) and *P. flexuosa* as the species with more dispersed foliage (longer petioles and low number of leaves) (Table 6). The other species remained in the intermediate region of the diagram and showed intermediate foliage structure.

**Fig. 5.**
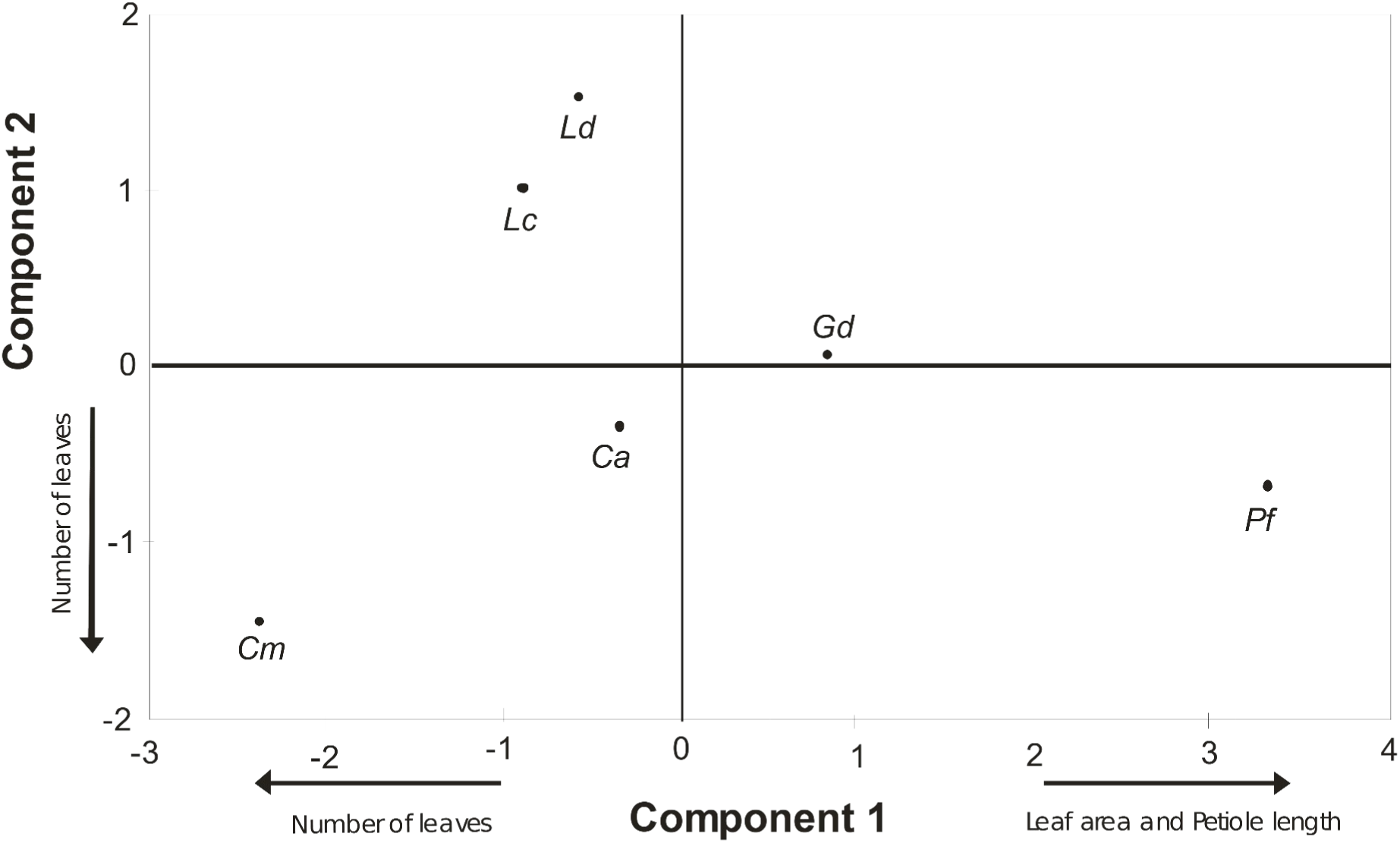
Position of woody plant species from Reserve of Ñacuñán along the first two axes obtained from a Principal Component Analysis carried out with structural foliage data. See Table 5 for loadings of variables for each component. See Table 1 for plant species codes. N= 10 for each plant species.

**Table 5.** Principal Component Analysis based on foliage structure variables of six woody plant species from Ñacuñán Reserve, Mendoza. Loadings for the most heavily weighted variables are shown in bold.

| *Variables* | *Code* | *Components* |  |  |
| --- | --- | --- | --- | --- |
|  |  | CP 1 | CP 2 | CP 3 |
| Large petiole | Pet | **0.84** | -0.28 | -0.16 |
| Number leaves above the plane | Leaf abov | -0.62 | 0.44 | 0.21 |
| Number leaves below the plane | Leaf low | **0.74** | -0.52 | 0.11 |
| Number leaves at the inner of branch | Num inn | -0.55 | **-0.63** | -0.19 |
| Number leaves at the middle of branch | Num mid | -0.7 | -0.54 | -0.36 |
| Number leaves at the exterior of branch | Num ext | **-0.73** | -0.34 | -0.26 |
| Area of leaf | Area | **0.88** | -0.24 | -0.15 |
| Branch distance to target branch (middle) | Branch mid | 0.26 | 0.38 | **-0.66** |
| Branch distance to target branch (exterior) | Branch ext | 0.08 | 0.46 | -0.59 |
| Eigenvalue |  | 3.82 | 1.77 | 1.13 |
| % Variance |  | 42.5 | 19.7 | 12.5 |
| Cumulative |  | 42.5 | 62.2 | **74.7** |

**Table 6:** Structural foliage variables (± SE) in six woody plants species at Ñacuñán Reserve. See Table 1 for plant species codes.

|  |  | Woody plant species |  |  |  |  |  |
| --- | --- | --- | --- | --- | --- | --- | --- |
|  |  | Pf | Ld | Gd | Ca | Cm | Lc |
| Petiole-length (mm) | | 21.1 ± 2.0 | 0.5 ± 0.0 | 5.2 ± 0.2 | 1.7 ± 0.1 | 0.4 ± 0.0 | 0.4 ± 0.0 |
| Leaf position (%) | *above* | 31 | 53.7 | 44.7 | 41 | 48.7 | 65.3 |
|  | *even with* | 21.3 | 32.3 | 21.7 | 29.3 | 31.3 | 14.3 |
|  | *below* | 46.7 | 14 | 33.7 | 29.7 | 20 | 20.3 |
| Number of leaves in 10 cm | *inner* | 4.7 ± 0.9 | 4.4 ± 3.3 | 11.7 ± 3.8 | 115.9 ± 33.5 | 233.4 ± 59.8 | 23.5 ± 10.2 |
| | *middle* | 17.1 ± 2.8 | 162.7 ± 29.4 | 22.1 ± 3.2 | 186.1 ± 31.7 | 579.5 ± 101.9 | 90.5 ± 27.5 |
| | *distal* | 20.5 ± 2.5 | 147.5 ± 21.7 | 32.7 ± 4.4 | 122.1 ± 32.1 | 521 ± 82.2 | 238.6 ± 45.8 |
| Leaf Area (cm<sup>2</sup>) | | 4.17 ± 0.2 | 0.37 ± 0.0 | 1.29 ± 0.1 | 0.32 ± 0.0 | 0.08 ± 0.0 | 0.23 ± 0.0 |
| Branch distance (middle) (cm) | | 13.5 ± 1.3 | 14.7 ± 1.4 | 10.3 ± 0.6 | 10.3 ± 0.4 | 9.9 ± 0.6 | 11.0 ± 1.1 |
| Branch distance (external) (cm) | | 13.9 ± 1.4 | 16.5 ± 1.3 | 9.6 ± 0.6 | 12.2 ± 1.3 | 12.0 ± 0.2 | 13.6 ± 1.4 |

## Discussion

The three bird species in the two habitats of Ñacuñán Reserve captured prey on trees and shrub foliage. The birds selected higher canopy heights (2 to 4.5 m) to realize their attacks in the mesquite woodlands and creosotebush shrubland although both habitats were dominated by foliage layers from 1 to 3 m (principally shrubs). The three bird species studied showed different patterns in the use of maneuvers to capture preys. The species with less flexibility in its spectrum of maneuvers was Ringed Warbling-Finch, who performed almost all its attacks by gleaning. The other two species were more flexible in their use of maneuvers, although the principal maneuver was not the same: Greater Wagtail-tyrant is primarily a gleaner species and Grey-crowned Tyrannulet is mainly a sally-hoverer species.

The three bird species showed a strong pattern of selection to *P. flexuosa* as feeding site in both habitat types studied (even in creosotebush shrubland where *P. flexuosa* coverage was only 6%, compared with 12-24% coverage in mesquite woodland). The birds species avoided the most abundant species: *L. divaricata* and *L. cuneifolia,* in mesquite woodland and creosotebush shrubland. Previous studies also found an important use of *P. flexuosa* by insectivorous birds in the Ñacuñán Reserve (Lopez de Casenave et al. 2008) and other sites within Monte Desert (Blendinger 2005).

Birds can modify their foraging behavior between habitats with different vegetation structure (Petit et al. 1990, Block 1990), however in this study the three bird species did not show differences in any of the foraging variables measured, despite the differences in foliage height profile and horizontal plants cover in mesquite woodland and creosotebush shrubland. The strong selection pattern of bird species for *P. flexuosa* determined that the height and substrate use were greatly influenced by this selection, and thus these foraging variables did not change between the three plots.

Holmes and Robinson (1981) found that gleaner birds have a stronger selection for woody plant species and are more influenced by foliage structure than hover birds. Under this scenario, dense and clustered foliage should be selected by the gleaner birds, due to this foliage facilitates the use of this maneuver; while open and not dense foliage, like *P. flexuosa*, would be less appropriate to perform that feeding maneuver. However, we did not find that insectivorous gleaner birds (Ringed Warbling-Finch and Greater Wagtail-tyrant) use vegetation in accordance with the foliage facilitation hypothesis, instead they selected sparse foliage of *P. flexuosa* to feeding. Regarding the hovered birds they might forage just as efficiently in plants with long petioles and leaves scattered apart as in plants with clustered leaves and short petioles and thus be less affected by differences in foliage structure (Holmes and Robinson 1981). However in our case, Grey-crowned Tyrannulet, had a slight tendency to sally-hover most frequently in *P. flexuosa* than in the others woody species. Bird species with a wide range of foraging techniques changed their techniques when foraging in tree species with different foliage structures (Unno 2002, Hino et al. 2002); in this study Grey-crowned Tyrannulet was the species with the wider maneuver spectrum, and thus this species could reduce constraints imposed by foliage structure, but this should be tested by any experiment or to compare attack rate in different plant species.

The other factor that could explain the bird selection is the prey abundance in the plants (Holmes and Robinson 1981, Holmes and Schultz 1988, Hino et al. 2002). We found that the most important variables for distinguished the plant species in terms of availability of arthropod were the greater abundance and biomass per individual plant and the diversity of orders and families of arthropod; these variables differed *P. flexuosa* from other plant species. By selecting this species, the three insectivorous bird species studied would maximize their search efficiency, since individuals of *P. flexuosa* are food patches with greater abundance and biomass of prey (by several orders of magnitude), so we believe that the size of each individual plant was a key factor determining the bird selection. The tree size may be an important factor to birds in several environments (Dean et al. 1999, Böhm et al. 2009), for example the large trees in desert areas may represent a keystone structure to avian assemblages (Seymour and Dean 2010, Thiele et al. 2008), increasing the bird diversity and abundance.

Another factor to considered is the nutritional content and secondary compounds in the plant foliage that can affect the richness, abundance and palatability of foliage arthropods and determine indirectly woody species selection by insectivorous birds (Recher et al. 1996, Greenberg and Bichier 2005). As in this research, the diversity of arthropods in *Larrea* sp. was low compared with other plant species of the central Monte desert (Debandi 1999), probably due to the large amounts of phenolic resin of their foliage (Rhoades 1977), to which only few arthropods are tolerant (Chapman et al. 1988). This chemical quality of foliage could explain the pattern of avoidance of *Larrea* spp. by birds. Contrary, *P. flexuosa* in our study had the greatest diversity in terms of the orders and families of arthropods and probably their foliage are more palatable to herbivorous insects (as the foliage of *Prosopis juliflora* in North American deserts, see Barth 1975, Maurer 1985). However detailed information about the bird diet is needed for confirming if the presence of certain arthropods is an important factor in the selection of feeding sites by birds.

In Ñacuñán Reserve, the richness and abundance of insectivorous birds both resident and migratory is greater in mesquite open woodland than in creosotebush shrubland during the breeding season (Cueto et al. 2006). There is general consensus that structural complexity of vegetation is a major driver of bird diversity (Wiens 1989); in Ñacuñán Reserve the birds could find more foraging sites in open woodland because there are more vegetation strata than in shrubland (Cueto et al 2006). In our study we can observed that these foraging sites are mainly provide by the *P. flexuosa* canopy. The same was observed in another place within the Monte (Telteca Reserve) where the resources associated with a single species of tree (*P. flexuosa*) appear to be as important for foraging birds (Blendinger 2005). The obtained data in the present research about the importance of *P. flexuosa* individuals, due to be an abundant food patch of great diversity of arthropod, provide important information about how birds use their habitat and which factors could be influencing the selection pattern of insectivorous birds of Monte desert.

## Appendix

Equations to estimated arthropod weight (W, mg) from body lengths measured (L, mm) of arthropods Orden or Families collected at Ñacuñán Reserve, central Monte desert, Argentina. For statistical analyses, we transformed the variable as W =ln W to reduce heteroscedasticity

-For Orden Araneae the dry weight estimate was:

W = 0.0497* L ^2.58^ (n = 10, r^2^= 0.99, P<0.0001)

-For Orden Coleoptera it was

W= 0.0428*L ^2.55^ (n = 9, r^2^ = 0.96, P<0.0001)

-For Heteroptera (without Family Pentatomidae), Homóptera, Orthoptera and Díptera (Suborden Brachycera)

W=0.0339*L^2.4^ (n = 19, r^2^= 0.95, P<0.0001)

-For Hymenoptera, Lepidoptera (adults), Solifuga and Diptera (Suborden Nematocera)

W= 0.0198*L^2.15^ (n = 12, r^2^= 0.9, P<0.0001)

-For Larvaes

W= 0.0174*L ^2.26^ (n = 10, r^2^= 0.95, P<0.0001)

-For Family Pentatomidae

W=0.0244*L^2.77^ (n = 6, r^2^= 0.96, P=0.0004)

## Notes

### Competing Interest Statement

The authors have declared no competing interest.

